# Fatty acid synthesis promotes inflammasome activation through NLRP3 palmitoylation

**DOI:** 10.1101/2023.10.30.564549

**Authors:** Stuart Leishman, Najd M. Aljadeed, Liyunhe Qian, Shamshad Cockcroft, Jacques Behmoaras, Paras K. Anand

**Affiliations:** Department of Infectious Disease, Imperial College London, London, W12 0NN UK; Department of Neuroscience, Physiology and Pharmacology, Division of Biosciences, University College London, London WC1E 6JJ, UK; Programme in Cardiovascular and Metabolic Disorders and Centre for Computational Biology, Duke-NUS Medical School Singapore, Singapore

**Keywords:** NLRP3, Inflammasome, Palmitoylation, FASN, Lipid metabolism

## Abstract

Inflammasomes are multi-protein complexes assembled by NOD-like receptor (NLR) family of proteins, which play critical roles in infectious, inflammatory and metabolic diseases. The assembly of the NLRP3 inflammasome is triggered upon recognition of an apt stimulus by the sensor protein, resulting in binding to pro-caspase-1 via the adaptor protein ASC. Inflammasome activation results in the maturation of the precursor forms of cytokines IL-1_β_ and IL-18, along with caspase-1-dependent pyroptosis, a pro-inflammatory form of cell death. Emerging evidence suggests the involvement of lipid metabolism in inflammasome activation; however, the precise mechanisms by which lipids regulate the NLRP3 inflammasome remain ambiguous. A multi-enzyme protein, fatty acid synthase (FASN) is a central regulator of lipid metabolism partaking an essential role in fatty acid biosynthesis pathway by catalysing the production of palmitic acid. Palmitic acid acts as a precursor to long-chain fatty acids and additionally regulates cellular functions by palmitoylation, a process in which palmitate is reversibly added to cysteine residues of target proteins, modifying protein localization and function. Here, we undertook a pharmacological approach to investigate the roles of fatty acid biosynthetic pathway in NLRP3 inflammasome activation. Our results demonstrated that inhibition of FASN in primary mouse and human macrophages abrogates the activation of the NLRP3 inflammasome, resulting in blunted caspase-1 activation. Furthermore, this phenomenon relied on protein palmitoylation as *in vitro* and *in vivo* abrogation of palmitoylation similarly reduced NLRP3 activation, which could be restored by exogenously supplementing palmitate in cultured cells. Consequently, an acyl biotin exchange assay corroborated NLRP3 palmitoylation. Notably, activation of the dsDNA sensing AIM2 inflammasome remained unaltered when either FASN or palmitoylation was blocked. These results therefore highlight the pivotal role of FASN and palmitoylation, shedding new mechanistic insights into the activation of the NLRP3 inflammasome.

## Introduction

The inflammasomes are multiprotein complexes which are assembled in response to infectious, endogenous, and environmental stimuli. The assembly of the inflammasome, composed of a sensor protein which is bound to pro-caspase-1 through the adaptor protein ASC, results in the cleavage of the inflammatory pro-caspase-1, which in turns results in the processing and maturation of cytokines pro-IL-1β and pro-IL-18 to their biologically active forms^1–4^. The mature caspase-1 also cleaves gasdermin-D (GSDMD) resulting in the N-terminal fragment of GSDMD translocating to the plasma membrane where it is involved in pore formation^5,6^. The pores thus formed allow cytokines and other cellular contents to be released extracellularly resulting in an inflammatory form of cell death called pyroptosis^5,6^. The well-studied NLRP3 inflammasome responds to a range of stimuli and comprises a two-step mechanism in which the NLRP3 is first expressed in an NF-kB dependent manner followed by the assembly of the inflammasome upon sensing apt pathogen-associated and danger-associated molecular patterns. Dysregulated NLRP3 inflammasome is implicated in a range of inflammatory and autoimmune diseases^7^. However, the ambiguity in the activation mechanisms of the NLRP3 inflammasome has impeded the development of precise therapeutics to target this inflammasome.

The activation mechanisms of the NLRP3 inflammasome are not completely understood. However, potassium efflux is largely accepted as a consensus event that must occur prior to NLRP3 inflammasome activation^8^. Moreover, post-translational modification of inflammasome components may further regulate NLRP3 inflammasome activity^9,10^. Emerging evidence over the past several years has implicated key roles for lipids in NLRP3 activation^11–13^. Notably, cholesterol egress from the endosome/lysosome compartment and subsequent trafficking to the ER is critical for the activation of the NLRP3 inflammasome^11^. While the detailed mechanism remains to be established, cholesterol may provide a conducive milieu in these organelles for efficient sensing by NLRP3. A role for cholesterol biosynthesis involving the transcription factor SREBP2 has also been reported to contribute to NLRP3 inflammasome activation^14^. On the contrary, the roles of SREBP1, the transcription factor that regulates the biosynthesis of fatty acid synthesis are less well defined. Fatty acid synthase (FASN), a major enzyme in the fatty acid biosynthesis pathway, has been implicated in immune signalling by regulating the TLR4 pathway and the levels of downstream inflammatory cytokine production^15,16^. This is largely attributed to the de novo activity of FASN fuelling sphingolipid production, thereby exerting influence on membrane raft formation and precise Rho-GTPase trafficking^15–18^. These studies therefore imply an indirect role for FASN in inflammasome priming. However, the role of FASN in NLRP3 inflammasome activation remains ambiguous.

Lipids have also been implicated in defining the precise localization of the NLRP3 inflammasome which has been debated almost since the discovery of the complex^19^. A recent study demonstrated that a polybasic region in the NLRP3 PYD to NACHT linker region associates with the phospholipid phosphatidylinositol 4-phosphate (PI4P) in the dispersed *trans* Golgi network (dTGN) before NLRP3 inflammasome can be assembled^12^. Other studies have demonstrated that the double-ring cage structure of the active NLRP3 inflammasome is promoted by membrane association^20^. These observations suggest that NLRP3 is tethered to a membrane. However, mechanisms necessary for the membrane association of the NLRP3 inflammasome remain ambiguous.

Despite gaining some understanding in recent years, the precise mechanism through which lipid metabolism influences NLRP3 activation remains ambiguous^21^. Additionally, a definite link between NLRP3 activation and lipid signalling remains elusive. In this study, we investigated the role of fatty acid biosynthesis pathway in inflammasome activation. Our data shows that the pharmacological blockade of FASN in mouse macrophages specifically regulates the activation of the NLRP3 inflammasome. This effect of FASN is intricately linked to NLRP3 palmitoylation. Consequently, ablation of cellular palmitoylation dampened NLRP3 inflammasome activation, which could be restored by exogenous addition of palmitate. By contrast, activation of the AIM2 inflammasome remained unrestricted. Moreover, these findings were replicated *in vivo* where the inhibition of FASN or palmitoylation led to a significant reduction in serum IL-1β levels. Altogether, these results demonstrate that fatty acid metabolism and NLRP3 palmitoylation play critical roles in the activation of the NLRP3 inflammasome.

## Results

### Inhibition of FASN reduces caspase-1 activation, IL-1*β* secretion, and pyroptosis

FASN is a crucial multi-functional enzyme involved in the de novo synthesis of fatty acids^22,23^. The enzyme utilizes acetyl-CoA and malonyl-CoA as substrates, converting them into saturated fatty acids in a series of reactions^22^. These fatty acids play key roles in structural integrity of the membranes, energy homeostasis, hormone production and as signalling mediators^16^. A previous study suggested a role for FASN in inflammasome priming by contributing to the production of lipids rafts which facilitate TLR4 signalling at the plasma membrane^15^. However, a direct role for fatty acid synthesis, and particularly FASN, in the activation of the NLRP3 inflammasome remains unknown. In order to interrogate this, we exposed LPS-primed mouse bone marrow-derived macrophages (BMDMs) to FASN inhibitor, cerulenin. Cerulenin is a natural mycotoxin which specifically and irreversibly inhibits the β-ketoacyl synthase activity resulting in a dead-end FAS inhibition^24^. It has also been demonstrated to exhibit selective cytotoxicity in several established cancer cell lines^25^. BMDMs exposed overnight to cerulenin displayed morphological features of cytotoxicity **(Fig. S1A)**. In order to specifically examine the role of FASN during NLRP3 activation, cerulenin was added to media 2h after LPS priming by which point the cells displayed abundant expression of NLRP3 sensor protein and additionally exhibited no characteristics of cytotoxicity as observed by light microscopy **(Fig. 1A, B)**. Subsequent activation of the NLRP3 inflammasome by ATP resulted in cleavage of pro-caspase-1 to its 20 kDa active form **(Fig. 1A)**. However, the processing of caspase-1 was significantly decreased in a dose-dependent manner in cells exposed to cerulenin demonstrating reduced NLRP3 inflammasome activation **(Fig. 1A, C)**. Active caspase-1 further leads to the cleavage of gasdermin D (GSDMD), the N-terminal of which translocates to the PM resulting in pore formation and induction of inflammatory cell death, pyroptosis^5,6^. In agreement with reduced caspase-1, inhibition of FASN resulted in a dose-dependent decrease in GSDMD N-terminal cleavage as compared to control cells **(Fig. 1A, C)**. Consequently, the secretion of caspase-1-dependent cytokine IL-1β also exhibited a significant decrease in response to cerulenin while the levels of secreted TNF-α and IL-6 remained unaltered **(Fig. 1D, E, Fig. S1B)**.

**Fig. 1.**
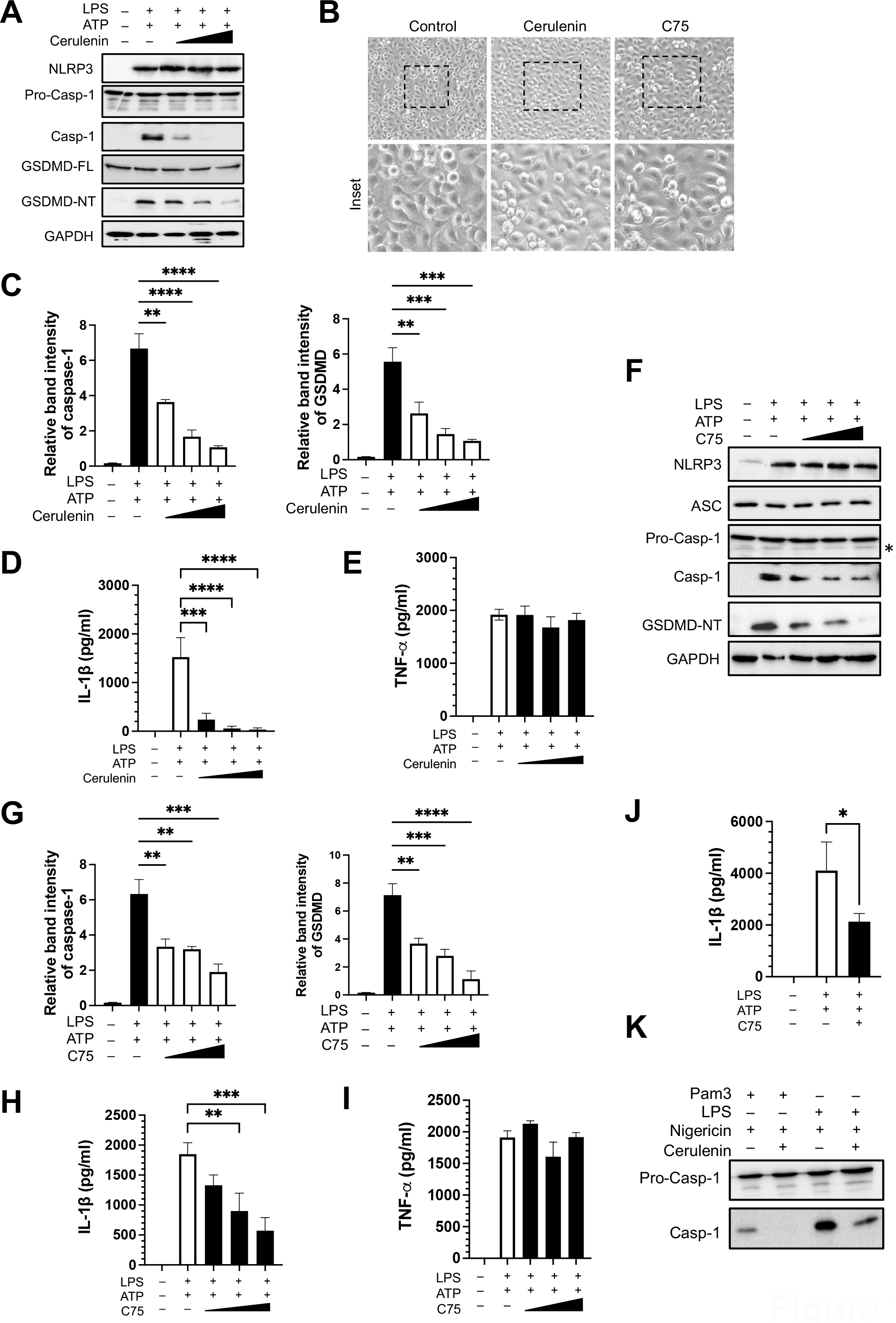
Inhibition of FASN reduces caspase-1 activation, IL-1β secretion, and pyroptosis. **(A)** BMDMs stimulated with inflammasome priming signal LPS (500 ng/ml; 3.5 hours) were exposed to increasing concentrations of the FASN inhibitor cerulenin (6.25, 12.5, and 25 μM; 1 hour) prior to exposure to ATP (5 mM; 45 min). Cell lysates were collected and immunoblotted with the indicated antibodies. GADPH was used as a loading control. **(B)** Light microscopy images of LPS-primed BMDMs treated either with cerulenin or C75, as above. Inset, magnification of the highlighted area. **(C)** Relative band intensity of cleaved caspase-1 and GSDMD-NT as measured by ImageJ and representative of the experiment above. **(D, E)** IL-1β and TNF-α released from cells treated as above. **(F)** BMDMs primed with LPS as above were exposed to FASN inhibitor C75 at increasing concentrations (12.5, 25, and 50 μM) 2 hours prior to stimulation with ATP. Cell lysates were collected and immunoblotted with the indicated antibodies. GADPH was used as a loading control. Asterisk (*) on the immunoblot denotes a non-specific band. **(G)** Relative band intensity of cleaved caspase-1 and GSDMD-NT as measured by ImageJ and representative of the experiment above. **(H, I)** IL-1β and TNF-α released from cells treated as above. **(J)** Primary human monocyte-derived macrophages were exposed to LPS and ATP in the absence or presence of C75. IL-1β release was measured in the supernatants by ELISA. **(K)** BMDMs were exposed to TLR2 agonist Pam3CKS4 (500 ng/ml; 3.5 hours) and nigericin (20 μM; 1 hour) or LPS and nigericin in the absence or presence of cerulenin. Cell lysates were immunoblotted with a caspase-1 antibody. Data shown are mean ± SD, and experiments shown are representative of at least three independent experiments. *, p<0.05; **, p < 0.01; ***, p < 0.001; ****, p < 0.0001, by Student’s *t* test.

In order to further validate these findings, we next employed a potent synthetic inhibitor of FASN, C75. C75 competitively interferes with the binding of malonyl-CoA to the β -ketoacyl synthase domain of FASN inhibiting long-chain fatty acid elongation^24^. Similar to the data obtained with cerulenin, exposure to C75 resulted in a decrease in caspase-1 activation and GSDMD processing compared to control cells **(Fig. 1F, G)**. In agreement, IL-1β secretion was also decreased in LPS-primed cells exposed to C75 even though the secretion of the inflammasome-independent cytokines TNF-α and IL-6 exhibited no change **(Fig. 1H, I, Fig. S1C)**. These results were also mirrored in primary human monocyte-derived macrophages where a lower concentration of C75 potently decreased IL-1β secretion **(Fig. 1J)**, and in BMDMs upon activation of the non-canonical inflammasome **(Fig. S1D)**. In order to further examine whether the response is specific to TLR4 or the purinergic receptor, P2X_7_R, we substituted stimuli and instead primed cells with TLR2 agonist, Pam3CSK4, and a microbial toxin, nigericin, as an activation signal in these experiments. In agreement with the results above, inhibition of FASN resulted in reduced caspase-1 activation suggesting that FASN is vital for NLRP3 inflammasome activation irrespective of the upstream priming and activation signals involved **(Fig. 1K)**. These results, therefore, demonstrate a critical role for fatty acid synthesis in calibrating the activation of the NLRP3 inflammasome.

### Palmitoylation is essential and specific to the activation of the NLRP3 inflammasome

The FASN pathway catalyzes the synthesis of a 16-carbon saturated fatty acid, palmitic acid, which serves as a precursor to long-chain fatty acids^26^. Additionally, palmitate can be directly utilised in palmitoylation (*S*-acylation), a post-translational modification in which palmitate is reversibly added to cysteine residues of target proteins enabling their association with cell membranes and influencing modulation of protein localization, function, and stability^27,28^. In order to next examine whether the FASN pathway regulates inflammasome through palmitoylation, we independently examined the impact of palmitoylation on the activation of the NLRP3 inflammasome. To probe this, we blocked palmitoylation by exposing cells to a competitive analogue of palmitate, 2-bromopalmitate (2-BP), and interrogated caspase-1 cleavage in response to NLRP3 activating stimuli. In parallel, a separate set of cells were exposed to cerulenin as a control. As expected, cerulenin diminished caspase-1 activation and cleavage of GSDMD N-terminal fragment **(Fig. 2A)**. In agreement, levels of the mature form of IL-1β were also decreased in response to cerulenin as compared to control cells **(Fig. 2A, C)**. Intriguingly, exposure to 2-BP resulted in a similar decrease in caspase-1 activation in a concentration dependent manner **(Fig. 2A, B)**. Similarly, the expression of GSDMD N-terminal and the cleaved IL-1β were also decreased **(Fig. 2A, B)**. Likewise, the secreted levels of IL-1β also exhibited a concentration-dependent decrease upon exposure to 2-BP as compared to control cells exposed to LPS+ATP **(Fig. 2C)**. However, the levels of secreted TNF-α and IL-6 remained unimpaired in response to either cerulenin or 2-BP **(Fig. 2D, Fig. S2)**.

**Fig. 2.**
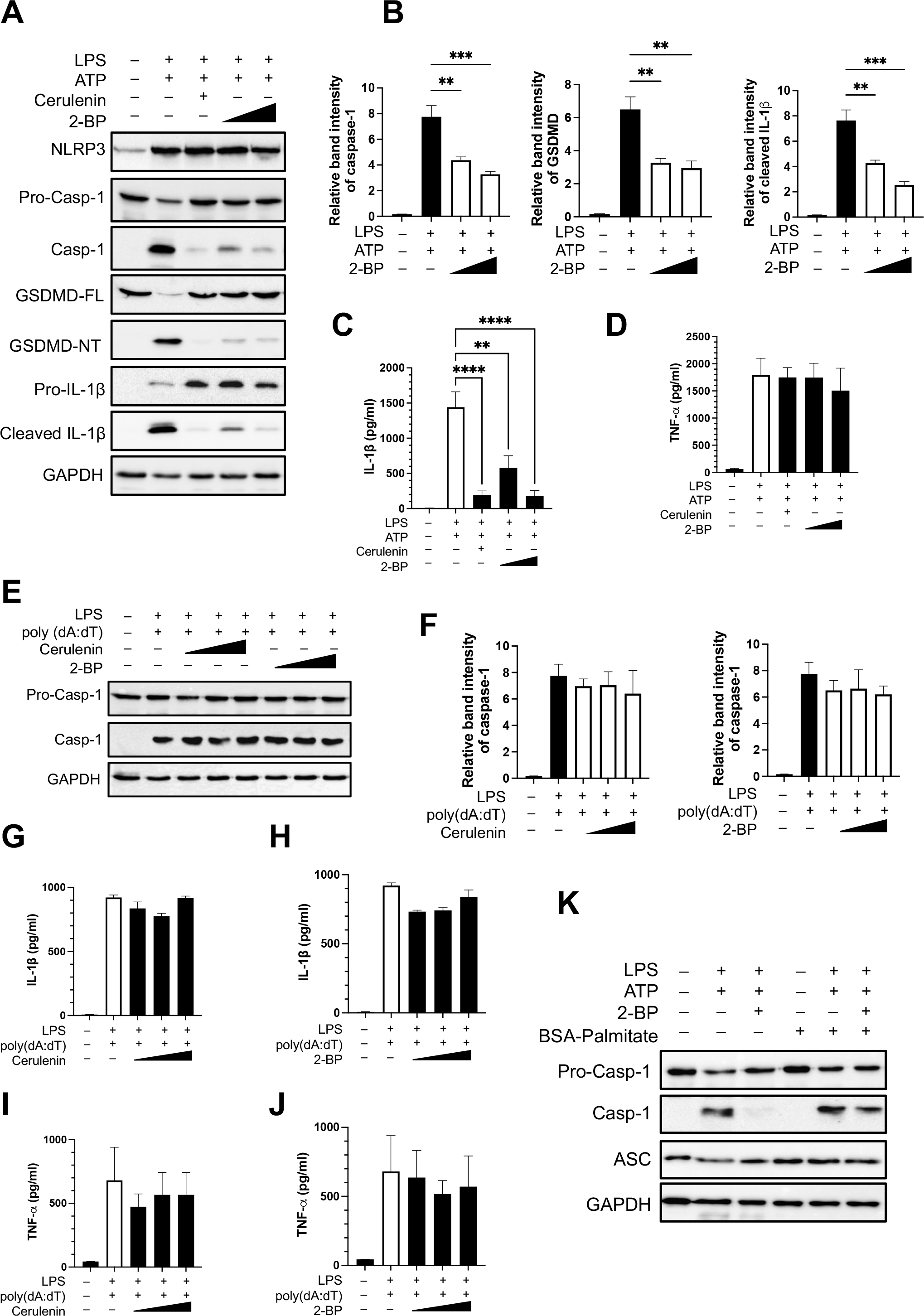
Palmitoylation is essential and specific to the activation of the NLRP3 inflammasome. **(A)** BMDMs exposed either to cerulenin (12.5 μM; 1 hour) or to increasing concentrations of the palmitoylation inhibitor, 2-BP (50 and 100 μM) and stimulated with NLRP3 agonist, LPS (500 ng/ml; 3.5 hours) and ATP (5 mM; 45 min). Cell lysates were collected and immunoblotted with the indicated antibodies. GADPH was used as a loading control. **(B)** Relative band intensity of cleaved caspase-1, GSDMD-NT, and cleaved IL-1β as measured by ImageJ and representative of the experiment above. **(C, D)** IL-1β and TNF-α released from cells treated as above. **(E)** BMDMs stimulated with LPS (500 ng/ml; 2 hours) were exposed either to increasing concentrations of cerulenin (6.25, 12.5, and 25 μM; 1 hour) or to increasing concentrations of the palmitoylation inhibitor, 2-BP (25, 50 and 100 μM) in cells transfected with AIM2 agonist, poly(dA:dT) overnight. Cell lysates were collected and immunoblotted with the indicated antibodies. GADPH was used as a loading control. **(F)** Relative band intensity of cleaved caspase-1 as measured by ImageJ and representative of the experiment above. **(G-J)** IL-1β and TNF-α released from cells treated as above. **(K)** BMDMs treated as in (A) above were additionally exposed to BSA-conjugated palmitate (500 μM) overnight in the indicated samples. Cell lysates were collected and immunoblotted with the indicated antibodies. GADPH was used as a loading control. Data shown are mean ± SD, and experiments shown are representative of at least three independent experiments. **, p < 0.01; ***, p < 0.001; ****, p < 0.0001, by Student’s *t* test.

Next, we examined whether the FASN/palmitoylation pathway broadly regulates inflammasome activation. In order to test this, cells were transfected with dsDNA to activate the DNA sensing AIM2 inflammasome in the presence of either cerulenin or 2-BP. Intriguingly, neither FASN nor palmitoylation regulated the AIM2 inflammasome **(Fig. 2E, F)**. Consequently, the expression of cleaved caspase-1 remained similar compared to control cells **(Fig. 2E, F)**. In agreement, the levels of secreted IL-1β, TNF-α, and IL-6 remained similar to that in control cells that were not exposed to any of the inhibitors **(Fig. 2G-J, Fig. S2)**.

Since 2-BP inhibited the NLRP3 inflammasome, we speculated that adding exogenous palmitate should rescue caspase-1 activation. Correspondingly, addition of BSA-conjugated palmitate to cells rescued the effect of 2-BP by elevating caspase-1 activation in macrophages exposed to NLRP3 stimuli **(Fig. 2K)**. Remarkably, while palmitate alone did not have an effect, control cells exposed to NLRP3 stimulus and supplemented with BSA-palmitate also displayed increased caspase-1 processing **(Fig. 2K)**. Altogether, these results suggest that palmitoylation is critical and specific to the activation of the NLRP3 inflammasome. Additionally, the results propose that palmitoylation can further augment NLRP3 inflammasome assembly, underscoring a regulatory role for this post-translational modification.

### NLRP3 is palmitoylated which is abrogated when FASN signalling is blocked

The dynamic control of palmitoylation is maintained by the activity of palmitoyl acyltransferases that add a palmitate on cysteine residues within the protein sequence^28^. Though the exact mechanisms remain unknown, the neighbouring amino acids and secondary structures can influence the specificity of palmitoylation^27,28^. In order to next decipher whether any of the inflammasome components is palmitoylated, we carried out *in-silico* approaches to predict putative palmitoylation sites in the amino acid sequence of mouse NLRP3, ASC and caspase-1 as described in detail in the methods section^29^. While the amino acid sequence of ASC and caspase-1 returned no putative sites, the mouse NLRP3 sequence exhibited six putative cysteine residues which can be palmitoylated.

Notably, five of the six sites were within the LRR domain of NLRP3 while one was found in the linker region between the NACHT and LRR domain **(Fig. S3)**.

In order to validate our *in-silico* studies and ascertain NLRP3 palmitoylation, we next established our findings by reconstituting the mouse NLRP3 inflammasome in HEK293T cells. After overnight transfection, HEK293T cells expressing the inflammasome components demonstrated caspase-1 cleavage which was reduced in the presence of both cerulenin and 2-BP demonstrating central roles of FASN and palmitoylation in inflammasome activation **(Fig. 3A)**. The regulation by FASN/palmitoylation was independent of whether the expressed proteins were human or mouse though HEK293T cells reconstituted with human NLRP3 inflammasome exhibited greater regulation by FASN and palmitoylation **(Fig. 3B)**. Overall, these data demonstrate that NLRP3 palmitoylation is fundamental to activation of the NLRP3 inflammasome and that this regulation is conserved between mice and humans.

**Fig. 3.**
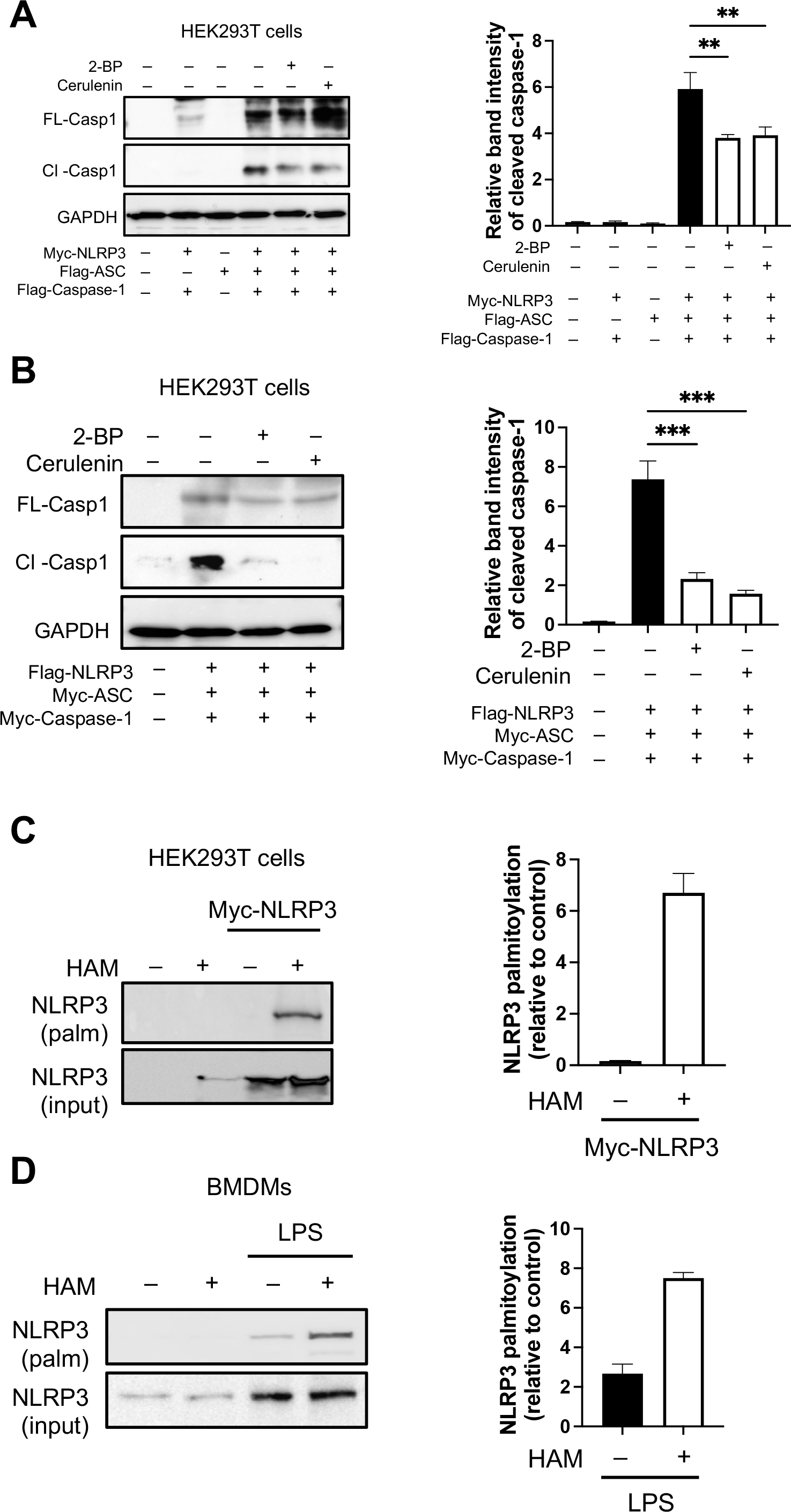
NLRP3 is palmitoylated which is abrogated when FASN signalling is blocked. NLRP3 inflammasome was reconstituted in HEK293T cells by transfecting with **(A)** either the mouse protein-coding plasmids, or **(B)** the human protein-coding plasmids. After 24 hours, cells were exposed to either cerulenin or 2-BP for 6 hours. Cell lysates were collected and immunoblotted for caspase-1. *Right panels*, Relative band intensity of cleaved caspase-1 as measured by ImageJ. **(C)** HEK293T cells were either transfected or not with wild-type Myc-NLRP3. After 24 hours, the cells were lysed, and the protein samples were divided into two parts. In HAM (+) samples, the palmitate residue was hydrolysed and exchanged with biotin for affinity enrichment while the HAM (-) sample served as the negative control. The levels of NLRP3 palmitoylation were detected by acyl-biotin exchange (ABE) assay by immunoblotting with anti-NLRP3 antibody. Total NLRP3 levels in the starting lysate served as the input control **(D)** BMDMs were exposed or not to LPS (500 ng/ml for 3.5 hours) and ABE assay was carried out to detect the levels of palmitoylated NLRP3. Total NLRP3 levels in the starting lysate served as the input control. Data shown are mean ± SD, and experiments shown are representative of at least three independent experiments. **, p < 0.01; ***, p < 0.001, by Student’s *t* test.

To next probe NLRP3 palmitoylation, we carried out an acyl-biotin exchange (ABE) assay in HEK293T cells transiently transfected overnight with Myc-tagged NLRP3. After 24h, cells were lysed followed by capping of the free cysteines in the lysates. The protein samples were divided into two fractions. In samples which are HAM (+), the palmitate residue was hydrolysed and exchanged with a thiol-reactive biotin moiety for affinity enrichment while the HAM (-) samples served as negative controls. Immunoblotting revealed the presence of *S*-acylated NLRP3 in HAM (+) HEK293T sample expressing Myc-NLRP3 while as expected HAM (-) and cells not expressing NLRP3 exhibited no acylation **(Fig. 3C)**. We further corroborated these findings in BMDMs validating palmitoylation of the endogenous NLRP3 in response to LPS treatment **(Fig. 3D)**. These studies therefore corroborate NLRP3 palmitoylation in line with our *in-silico* data and define palmitoylation as a key mechanism to regulate NLRP3 activity.

### FASN/palmitoylation axis is important for inflammasome assembly in vitro and in vivo

We next sought to determine the mechanism by which palmitoylation modified NLRP3 activation. As protein palmitoylation is typically coupled to membrane association influencing protein function, we hypothesized that palmitoylation might control the proximal event of NLRP3 assembly. Activation of the inflammasome and subsequent IL-1β secretion is dependent on the formation of a supramolecular complex termed ASC ‘specks’, which are found in detergent-insoluble cellular fraction^30^. As expected, stimulation of BMDMs with NLRP3 agonist resulted in the formation of a 1 μm ASC ‘speck’ which was visible in control cells by confocal microscopy **(Fig. 4A)**. However, ASC specks were significantly abolished in cells exposed to either cerulenin or 2-BP under conditions that activated the NLRP3 inflammasome **(Fig. 4A, B)**. We next independently interrogated whether AIM2 inflammasome assembly is regulated by FASN or palmitoylation. To probe this, mouse macrophages were transfected with dsDNA, which resulted in increased ASC speck formation. However, as expected, ASC speck formation was not restricted when either FASN or palmitoylation were pharmacologically blocked **(Fig. 4C, D)**. Furthermore, we considered the extent of ASC oligomerisation by cross-linking Triton X-100-insoluble fraction with disuccinimidyl suberate^30^. This treatment yielded most of the ASC as highly oligomerised in ATP treated control cells but not in cells exposed to cerulenin or 2-BP **(Fig. 4E)**. Taken together, these data propose that palmitoylation is pivotal to the assembly of the NLRP3 inflammasome.

**Fig. 4.**
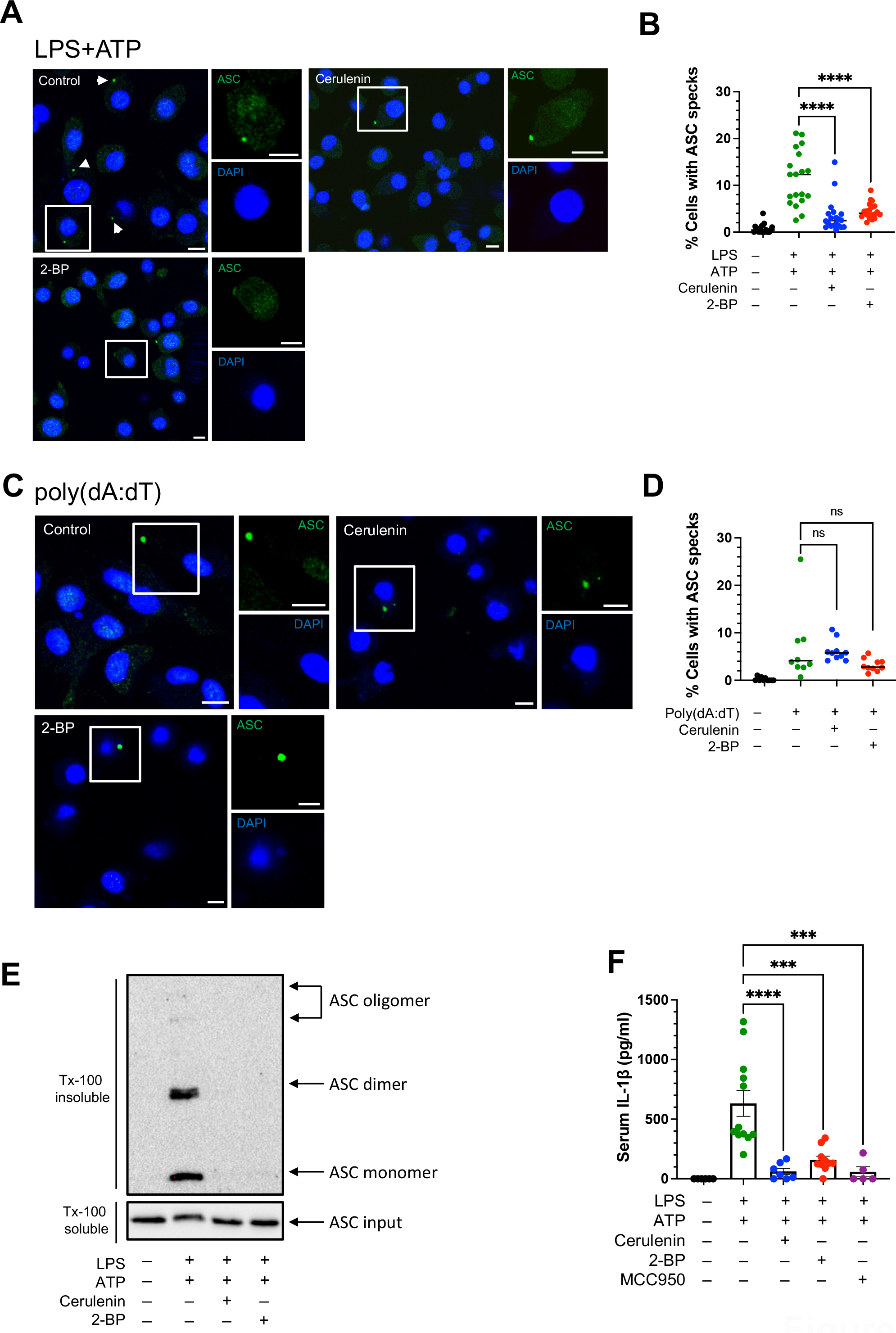
FASN/palmitoylation axis is important for inflammasome assembly *in vitro* and *in vivo*. **(A)** BMDMs were exposed to LPS+ATP in the absence or presence of cerulenin (12.5 μM) or 2-BP (100 μM). Cells were fixed and labelled with anti-ASC antibody to visualize ASC specks. DNA was stained with DAPI. **(B)** Quantitative analysis of the percentage of cells with ASC specks in samples treated as above. Each dot represents an individual field with at least n=30 cells. **(C)** LPS-primed BMDMs transfected with poly(dA:dT) to activate the AIM2 inflammasome were exposed or not to cerulenin (12.5 μM) or 2-BP (100 μM). Cells were fixed and labelled with anti-ASC antibody to visualize ASC specks. DNA was stained with DAPI. **(D)** Quantitative analysis of the percentage of cells with ASC specks in samples treated as above. Each dot represents an individual field with at least n=30 cells. **(E)** BMDMs were exposed to LPS+ATP in the absence or presence of cerulenin (12.5 μM) or 2-BP (100 μM). ASC oligomerization in Triton x-100 insoluble fraction was analysed by immunoblotting after cross-linking with DSS. The initial lysate immunoblotted with anti-ASC antibody served as an input control. **(F)** C57BL/6 mice were administered intraperitoneally (i.p.) with vehicle control, or cerulenin (50 mg kg^-1^), or 2-BP (100 mg kg^-1^), or MCC950 (50 mg kg^-1^), and LPS (100 μg kg^-1^). After 4 hours, mice were i.p. injected with ATP for 15 min. IL-1β was detected in the serum by ELISA. Data shown are mean ± SEM, and experiments shown are representative of at least three independent experiments. Scale bars, 5 μm. n.s., not significant. ***, p < 0.001; ****, p < 0.0001, by one-way Anova.

We next used a murine peritonitis model to investigate the role of FASN and palmitoylation pathway *in vivo*. To examine this, C57BL6/J mice were injected intraperitoneally with either FASN inhibitor cerulenin, or palmitoylation inhibitor 2-BP, or NLRP3 inhibitor MCC950 at the same time as LPS. After 4 hours, the NLRP3 inflammasome was activated by intraperitoneal administration of ATP, and IL-1β release was measured in the serum by ELISA. ATP administration induced a significant increase in serum IL-1β levels and this was considerably diminished by NLRP3 inhibitor, MCC950 **(Fig. 4F)**. Intriguingly, *in vivo* abolition of FASN or palmitoylation displayed a decrease similar to that observed with MCC950 indicating an NLRP3-dependent response **(Fig. 4F)**. These data show that FASN/palmitoylation axis is important for NLRP3 inflammasome activation both *in vitro* and *in vivo*.

## Discussion

Recent studies have demonstrated a correlation between metabolic lipid disorders and the activation of the NLRP3 inflammasome, directly implicating lipids as a potential contributor to the development of inflammatory diseases. However, the precise mechanism linking lipids to the activation of the NLRP3 inflammasome remains unclear. Here, we demonstrate that fatty acid biosynthesis pathway is central to activation of the NLRP3 inflammasome. Pharmacological blockade of a critical enzyme in the pathway, FASN, resulted in abrogation of caspase-1 activation and IL-1β secretion in response to NLRP3 activation. Notably, these findings were dependent on palmitoylation, a post-translational mechanism in which palmitic acid is added to the cysteine residue of target proteins influencing their localization and function. In agreement, we observed that NLRP3 is palmitoylated and blocking palmitoylation blunted inflammasome signalling which could be restored in the presence of exogenous palmitate.

A previous study examining the role of FASN revealed a key role for the enzyme in the production of inflammatory cytokines including TNF-α, IL-6, and IL-1β in response to TLR4 ligation^15^. Upon pharmacological inhibition of FASN, it was observed that the subsequent LPS-dependent signalling is compromised^15^. This was attributed to the limited production of acetoacetyl-CoA which contributed to the formation of lipid rafts that enable TLR4 signalling^15^. Other studies have confirmed a role for FASN in inflammatory signalling^31^. Our study by specifically investigating the role of FASN following TLR4 ligation highlights that FASN engages unique mechanisms to regulate inflammatory signalling and inflammasome activation.

Two recent studies have demonstrated that palmitoylation of GSDMD is important for pyroptosis induction and thus IL-1β secretion^31,32^. These studies identified that GSDMD-NT palmitoylation at Cys191/192 (human/mouse) position regulated membrane translocation of the N-terminal fragment although the prior processing of the GSDMD protein remained unaltered. Remarkably, our experiments revealed an upstream role for palmitoylation in NLRP3 inflammasome assembly. Accordingly, we observed palmitoylation of NLRP3 both in HEK293T cells transiently expressing NLRP3 and the endogenous NLRP3 in primary BMDMs by ABE assay. Consequently, both caspase-1 activation and IL-1β processing were blunted in the presence of a palmitoylation inhibitor. These results therefore shed light on previously elusive mechanisms through which palmitoylation regulates distinct steps in inflammasome assembly and signalling.

A very recent study identified zDHHC12 as the acyltransferase that adds palmitate at Cys844 position in NLRP3^33^. However, animals lacking zDHHC12 exhibited increased severity during an endotoxin shock model. In agreement, *Zdhhc12*^*-/-*^ macrophages also revealed increased IL-1β secretion. It was therefore concluded that palmitoylation dampens inflammasome activation. In our study, we observed that palmitoylation promotes caspase-1 activation and IL-1β secretion both in WT macrophages and in reconstituted HEK293T cells exposed to a well-established palmitoylation inhibitor, 2-BP. Moreover, the inhibitory effect of 2-BP could be rescued by supplementing cells with palmitic acid. This depicts differences in results possibly due to the different approaches used in the two studies. In the previous study, assays demonstrating caspase-1 activation and IL-1β secretion were only carried out in *Zdhhc12*^*-/-*^ macrophages. Since approximately 10% of the proteins coded by the human genome is palmitoylated^28,34^, the nature and identity of all zDHHC12 targets remain unknown.

Palmitoylation typically defines the membrane association of proteins influencing their localization and function. Therefore, our study suggests that NLRP3 or the inflammasome complex is tethered to an organelle. One study demonstrated that a polybasic region (residues 127-146) promotes association of NLRP3 with PI4P accumulated in the dTGN^12^. Notably, the dispersal of TGN38-postive vesicles, which have since been suggested to be endosomal in nature, occurred prior to NLRP3 puncta formation suggesting that PI4P accumulation is a prerequisite for NLRP3 activation^12^. Moreover, cryo-EM approaches independently revealed that the membrane association promotes NLRP3 inflammasome assembly^20^. Together with our studies, this suggests that both NLRP3 palmitoylation and association with negatively charged PI4P on dTGN may be required for NLRP3 inflammasome activation. Altogether, our study identifies NLRP3 palmitoylation as a regulatory mechanism controlling inflammasome activation, providing a novel target for calibrating NLRP3 activity in infectious and inflammatory diseases.

## Acknowledgements

This work was in-part supported by a grant from The Medical Research Council, UK (MR/S00968X/1) to P.K.A. The funders had no role in the design, conduct, or preparation of this manuscript.

## Author Contributions

S.L. and N.A. contributed to experimental design; S.L, N.A., L.Q. performed experiments; S.L, N.A., and P.K.A. analyzed data; S.C. and J.B. provided intellectual input and participated in discussions, P.K.A. designed the study, provided resources, overall supervision and wrote the original draft; all authors approved the final version.

## References

1. Swanson, K. V., Deng, M. & Ting, J. P. Y. The NLRP3 inflammasome: molecular activation and regulation to therapeutics. Nature Reviews Immunology vol. 19 477–489 Preprint at 10.1038/s41577-019-0165-0 (2019).

2. Man, S. M. & Kanneganti, T. D. Regulation of inflammasome activation. Immunol Rev 265, 6–21 (2015).

3. Anand, P. K., Malireddi, R. K. S. & Kanneganti, T.-D. Role of the Nlrp3 inflammasome in microbial infection. Front Microbiol 2, (2011).

4. Kanneganti, T.-D. et al. Bacterial RNA and small antiviral compounds activate caspase-1 through cryopyrin/Nalp3. Nature 440, 233–236 (2006).

5. He, W. T. et al. Gasdermin D is an executor of pyroptosis and required for interleukin-1_β_secretion. Cell Res 25, 1285–1298 (2015).

6. Liu, X. et al. Inflammasome-activated gasdermin D causes pyroptosis by forming membrane pores. Nature 535, 153–158 (2016).

7. Mangan, M. S. J. et al. Targeting the NLRP3 inflammasome in inflammatory diseases. Nat Rev Drug Discov 17, 588–606 (2018).

8. Muñoz-Planillo, R. et al. K+ Efflux Is the Common Trigger of NLRP3 Inflammasome Activation by Bacterial Toxins and Particulate Matter. Immunity 38, 1142–1153 (2013).

9. Song, N. et al. NLRP3 Phosphorylation Is an Essential Priming Event for Inflammasome Activation. Mol Cell 68, 185–197.e6 (2017).

10. Niu, T. et al. NLRP3 phosphorylation in its LRR domain critically regulates inflammasome assembly. Nature Communications 2021 12:1 12, 1–15 (2021).

11. de la Roche, M. et al. Trafficking of cholesterol to the ER is required for NLRP3 inflammasome activation. Journal of Cell Biology 217, 3560–3576 (2018).

12. Chen, J. & Chen, Z. J. PtdIns4P on dispersed trans-Golgi network mediates NLRP3 inflammasome activation. Nature 564, 71–76 (2018).

13. Zhang, Z. et al. Protein kinase D at the Golgi controls NLRP3 inflammasome activation. Journal of Experimental Medicine 214, (2017).

14. Guo, C. et al. Cholesterol Homeostatic Regulator SCAP-SREBP2 Integrates NLRP3 Inflammasome Activation and Cholesterol Biosynthetic Signaling in Macrophages. Immunity 49, 842–856.e7 (2018).

15. Carroll, R. G. et al. An unexpected link between fatty acid synthase and cholesterol synthesis in proinflammatory macrophage activation. Journal of Biological Chemistry 293, 5509–5521 (2018).

16. Wei, X. et al. Fatty acid synthesis configures the plasma membrane for inflammation in diabetes. Nature 539, 294–298 (2016).

17. Olona, A. et al. Sphingolipid metabolism during Toll-like receptor 4 (TLR4)-mediated macrophage activation. Br J Pharmacol 178, 4575–4587 (2021).

18. Muralidharan, S. et al. Immunolipidomics Reveals a Globoside Network During the Resolution of Pro-Inflammatory Response in Human Macrophages. Front Immunol 13, 926220 (2022).

19. Hamilton, C. & Anand, P. K. Right place, right time: Localisation and assembly of the nlrp3 inflammasome. F1000Res 8, (2019).

20. Andreeva, L. et al. NLRP3 cages revealed by full-length mouse NLRP3 structure control pathway activation. Cell 184, 6299–6312.e22 (2021).

21. Olona, A., Leishman, S. & Anand, P. K. The NLRP3 inflammasome: regulation by metabolic signals. Trends Immunol 43, 978–989 (2022).

22. Yaqoob, P. Fatty acids as gatekeepers of immune cell regulation. Trends in Immunology vol. 24 639–645 Preprint at 10.1016/j.it.2003.10.002 (2003).

23. Jensen-Urstad, A. P. L. & Semenkovich, C. F. Fatty acid synthase and liver triglyceride metabolism: housekeeper or messenger? Biochim Biophys Acta 1821, 747 (2012).

24. Flavin, R., Peluso, S., Nguyen, P. L. & Loda, M. Fatty acid synthase as a potential therapeutic target in cancer. Future Oncol 6, 551 (2010).

25. Kuhajda, F. P. Synthesis and antitumor activity of an inhibitor of fatty acid synthase. Proceedings of the National Academy of Sciences 97, 3450–3454 (2000).

26. Koundouros, N. & Poulogiannis, G. Reprogramming of fatty acid metabolism in cancer. British Journal of Cancer 2019 122:1 122, 4–22 (2019).

27. Linder, M. E. & Deschenes, R. J. Palmitoylation: policing protein stability and traffic. Nature Reviews Molecular Cell Biology 2007 8:1 8, 74–84 (2007).

28. Tabaczar, S., Czogalla, A., Podkalicka, J., Biernatowska, A. & Sikorski, A. F. Protein palmitoylation: Palmitoyltransferases and their specificity. Exp Biol Med 242, 1150 (2017).

29. Ning, W. et al. GPS-Palm: a deep learning-based graphic presentation system for the prediction of S-palmitoylation sites in proteins. Brief Bioinform 22, 1836–1847 (2021).

30. Fernandes-Alnemri, T. et al. The pyroptosome: a supramolecular assembly of ASC dimers mediating inflammatory cell death via caspase-1 activation. Cell Death Differ 14, 1590 (2007).

31. Balasubramanian, A. et al. Palmitoylation of gasdermin D directs its membrane translocation and pore formation in pyroptosis. bioRxiv (2023) doi:10.1101/2023.02.21.529402.

32. Du, G. et al. ROS-dependent palmitoylation is an obligate licensing modification for GSDMD pore formation. bioRxiv (2023) doi:10.1101/2023.03.07.531538.

33. Wang, L. et al. Palmitoylation prevents sustained inflammation by limiting NLRP3 inflammasome activation through chaperone-mediated autophagy. Mol Cell 83, 281–297.e10 (2023).

34. Sanders, S. S. et al. Curation of the Mammalian Palmitoylome Indicates a Pivotal Role for Palmitoylation in Diseases and Disorders of the Nervous System and Cancers. PLoS Comput Biol 11, 1004405 (2015).

