## Supplementary material for "Fatty acid synthesis promotes inflammasome activation through NLRP3 palmitoylation": Fig. S

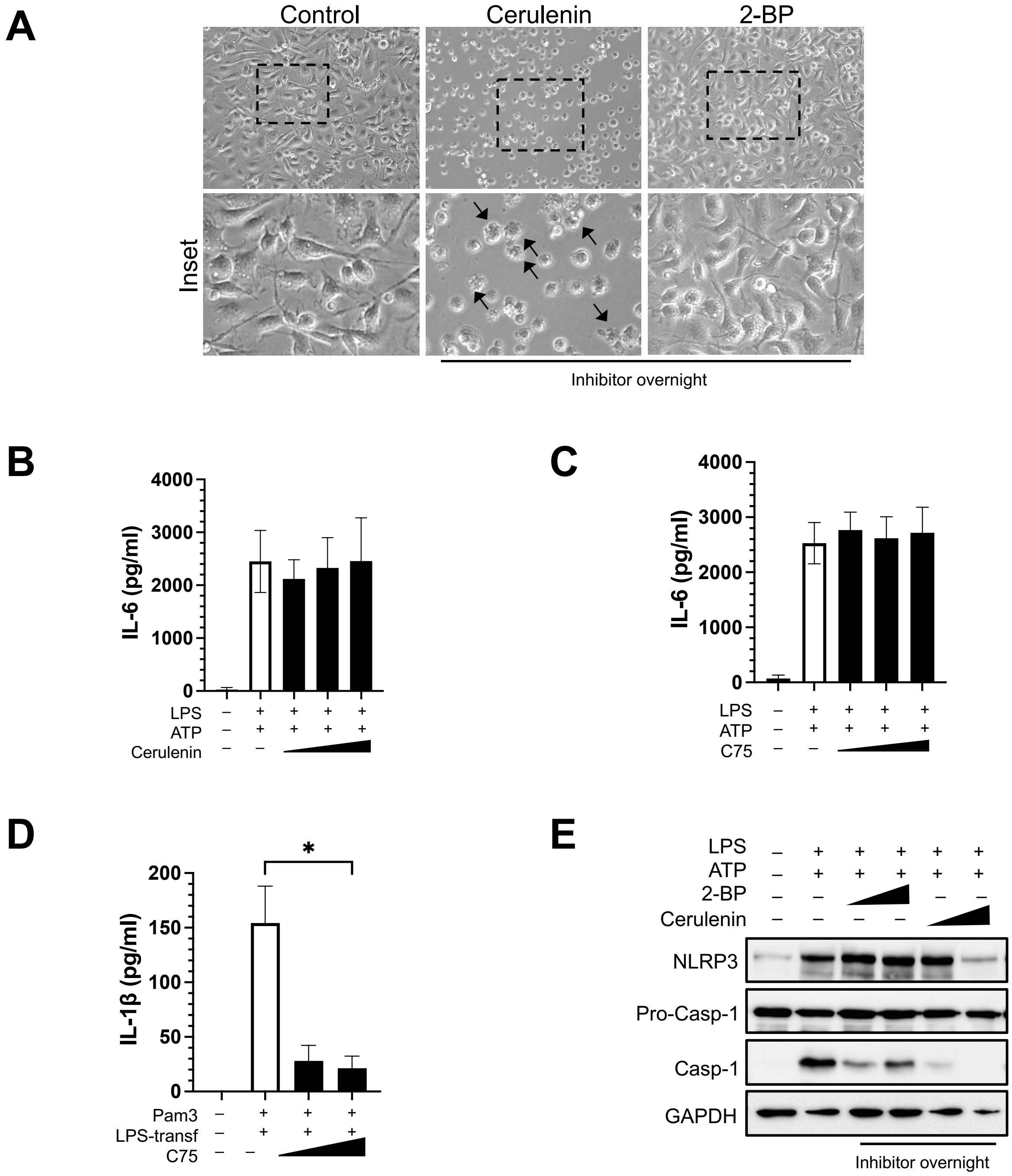


**Supplementary Fig. 1 (A)** BMDMs were exposed to FASN inhibitor cerulenin or palmitoylation inhibitor 2-BP overnight. Images were acquired on a light microscope. Inset, magnification of the highlighted area. **(B, C)** BMDMs stimulated with inflammasome priming signal LPS (500 ng/ml; 3.5 hours) were exposed to increasing concentrations of the FASN inhibitor cerulenin (6.25, 12.5, and 25 μM; 1 hour), or C75 (12.5, 25, and 50 μM; 2 hours) prior to exposure to ATP (5 mM; 45 min). IL-6 levels were measured in the cell supernatant by ELISA. **(D)** BMDMs were exposed to Pam3CSK4 (500 ng/ml; 4 hours) before transfecting with 1 μg/ml LPS overnight. At the end of the treatment, supernatants were assayed for IL-1β release by ELISA. **(E)** BMDMs were treated as in (A) above. Cell lysates were collected and immunoblotted with the indicated antibodies. GADPH was used as a loading control. Data shown are mean ± SD, and experiments shown are representative of at least three independent experiments. *, $p < 0.05$, by Student's *t* test.

### Figure S1

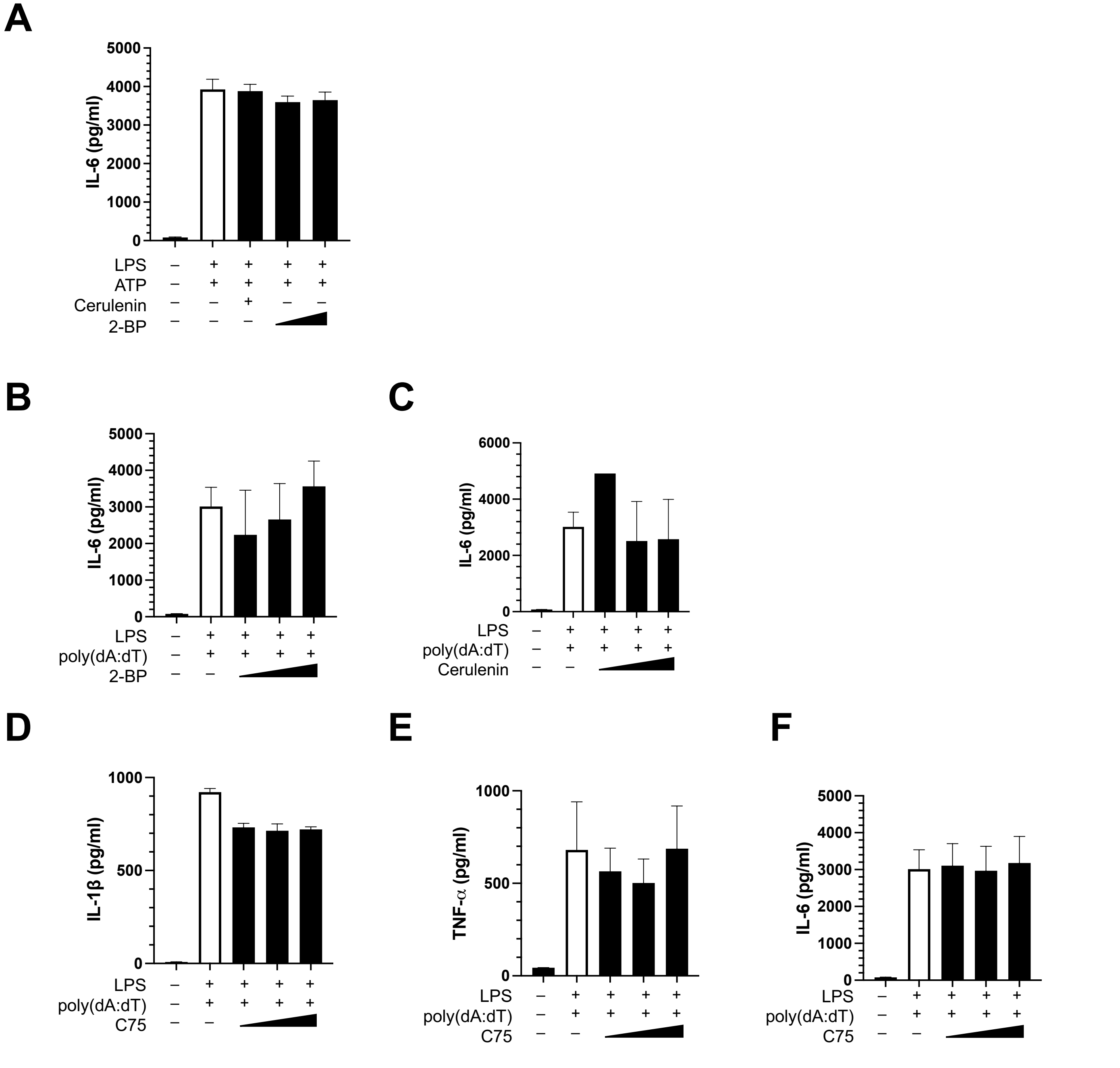


**Supplementary Fig. 2 (A)** BMDMs exposed either to cerulenin (12.5 µM; 1 hour) or to increasing concentrations of the palmitoylation inhibitor, 2-BP (50 and 100 µM) and stimulated with NLRP3 agonist, LPS (500 ng/ml; 3.5 hours) and ATP (5 mM; 45 min). At the end of the treatments, supernatants were collected and assayed for IL-6 release by ELISA. **(B, C)** BMDMs stimulated with LPS (500 ng/ml; 2 hour) were exposed either to increasing concentrations of cerulenin (12.5 µM; 1 hour) or to increasing concentrations of the palmitoylation inhibitor, 2-BP (25, 50 and 100 µM) in cells exposed to the AIM2 agonist, poly(dA:dT) overnight. At the end of the treatments, supernatants were collected and assayed for IL-6 release by ELISA. **(D-F)** BMDMs stimulated with LPS (500 ng/ml; 2 hour) were exposed to FASN inhibitor C75 at increasing concentrations (12.5, 25, and 50 µM), and further transfected with the AIM2 agonist, poly(dA:dT) overnight. At the end of the treatments, supernatants were collected and assayed for IL-1β, TNF-α, and IL-6 release by ELISA. Data shown are mean ± SD, and experiments shown are representative of at least three independent experiments.

### Figure S2

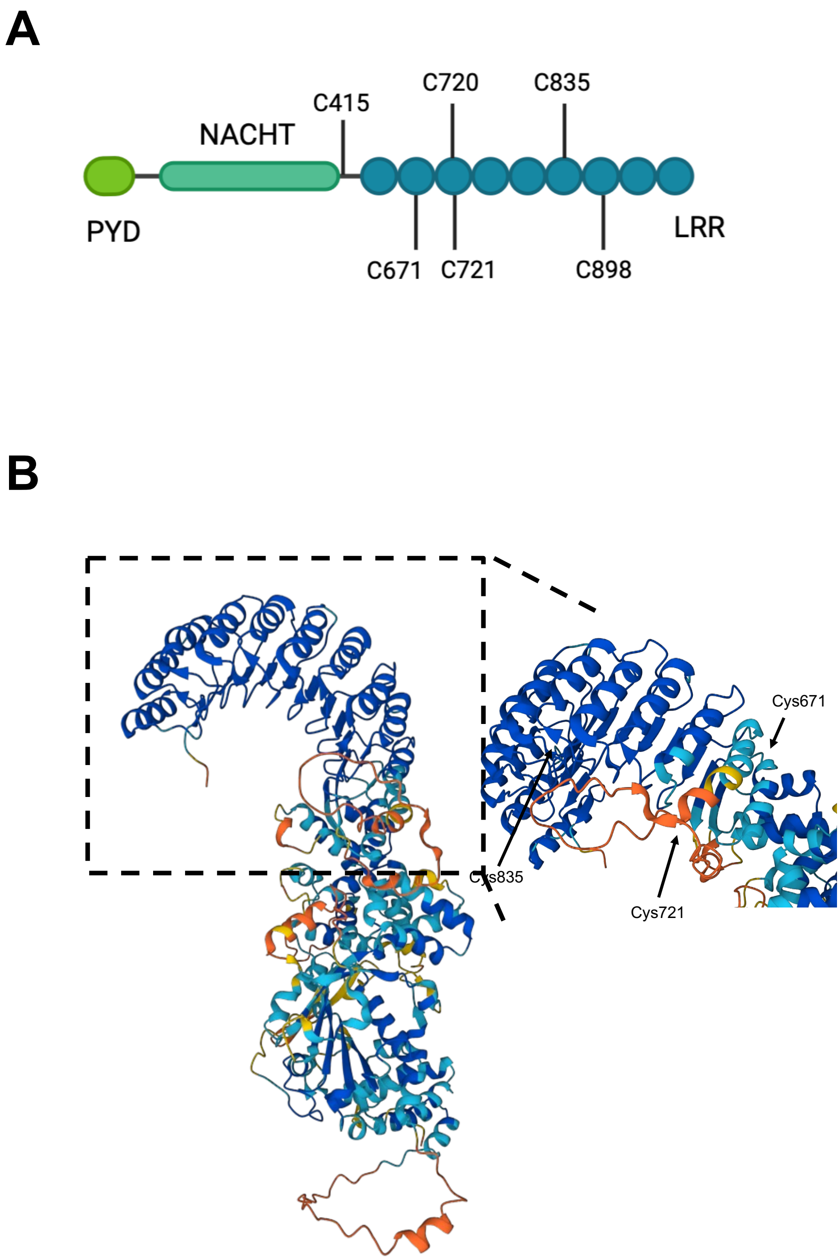


**Supplementary Fig. 3 (A)** Schematic depicting the palmitoylation sites identified in the mouse NLRP3 amino acid sequence by GPS-Palm. **(B)** Mouse NLRP3 AlphaFold protein structure depicting some of the putative Cys residues that can be palmitoylated in the LRR domain of the protein.

### Figure S3
